# XPO1 Inhibition by Selinexor Induces Nuclear p53 and p21 Accumulation, Cell-Cycle Arrest and Apoptosis in Testicular Germ Cell Tumors

**DOI:** 10.64898/2026.09.14.750914

**Authors:** Rashidul Islam, Anne Waldhaus, Isabella Federica Bollen, Lea Descher, Amir Shalan, Valentina Alejandra Araya Balde, Annalena Liesen, Andjela Kovacevic, Gina Esther Merges, Glen Kristiansen, Hubert Schorle

## Abstract

**Background:** Testicular germ cell tumors (TGCTs) are the most common solid malignancies in young men. Despite high cure rates with cisplatin-based chemotherapy, resistance and long-term toxicities remain clinical challenges in TGCTs. Selinexor, an FDA-approved XPO1 inhibitor, has demonstrated anti-tumor activity in other cancers, but its potential in TGCTs remains unknown.

**Objective:** We investigated the antitumor effects of selinexor in TGCT cell lines.

**Materials and methods:** XPO1 RNA expression, protein abundance and localization were evaluated by immunohistochemistry in TGCT tissue microarrays and non-neoplastic testicular tissues, and by meta-analysis of publicly available gene-expression microarray datasets from normal testis, primary TGCT specimens as well as TGCT cell lines. Selinexor effects on cell viability, cell-cycle distribution and apoptosis were evaluated using XTT assay and flow cytometry. p53 and p21 expression and localization were analyzed by Western blotting and Immunofluorescence.

**Results:** XPO1 expression was heterogeneous across normal testis, TGCT tissues and cell lines. Yet, selinexor reduced TGCT cell viability and induced G1 or G2/M cell-cycle arrest, apoptosis, increased total p53 and p21 protein levels, and their nuclear accumulation. Of note, control fibroblast exhibited limited sensitivity to selinexor.

**Discussion and conclusion:** Our findings demonstrate that inhibition of XPO1 with selinexor might be potential therapeutic strategy for TGCTs.

## 1. INTRODUCTION

Testicular cancer is the most common solid malignancy in men aged 15-44 years, and its incidence has increased worldwide over recent decades (1–3). Approximately 90-95% of testicular cancers are testicular germ cell tumors (TGCTs), which are broadly classified as seminomas and non-seminomas (1–3). Non-seminomatous germ cell tumors comprise embryonal carcinoma, choriocarcinoma, yolk-sac tumor, and teratoma (1–3). Standard treatment for TGCTs comprises radical inguinal orchiectomy followed by surveillance, retroperitoneal lymph-node dissection, radiotherapy or cisplatin-based chemotherapy, depending on tumor histology and disease stage (1–3). Nevertheless, despite the high curability of TGCTs, 3-5% of patients overall and 10-15% of TGCTs with metastasis develop cisplatin resistance, resulting in relapsed or refractory disease for which therapeutic options remain limited (1, 2, 4). Besides, conventional treatments can cause substantial long-term toxicities, including pulmonary, renal, neurological and ototoxic complications; impaired fertility and hypogonadism; cardiovascular morbidity; secondary malignancies; and persistent psychological distress (1, 2, 4). Together, these limitations highlight the need for alternative therapeutic strategies that improve treatment outcomes while reducing the overall treatment burden.

Exportin 1 (XPO1; also known as chromosome region maintenance 1, CRM1) is a karyopherin-β nuclear export receptor that maintains cellular homeostasis by regulating the nucleocytoplasmic localization of numerous protein and RNA cargos (5–8). In cancer, XPO1 dysregulation alters the localization of cargos involved in cell-cycle control, stress responses, and tumor progression (5–8). Overexpression XPO1 expression has been reported in multiple myeloma (9, 10), acute myeloid leukemia (11, 12), diffuse large B-cell lymphoma (13), mantle cell lymphoma (14) and chronic lymphocytic leukemia (15), lung (16), pancreatic (17), esophageal (18), colorectal (19), gastric cancers (20, 21). XPO1 overexpression at both the mRNA and protein levels has been associated with tumor progression, therapeutic resistance and poor clinical outcomes across multiple malignancies (5–9, 11, 13, 22–28).

Studies revealed that inhibiting XPO1 represents an attractive therapeutic strategy in cancer by preventing nuclear export of tumor-suppressor proteins and cell-cycle regulators, thereby conserving their nuclear functions and enforcing cell cycle arrest (5–8, 22, 23, 29, 30). In parallel, XPO1 block can attenuate the cytoplasmic translation of oncogenic mRNAs and compromise DNA-damage-repair capacity, collectively promoting apoptotic cell death (5–8, 22, 25, 29, 31, 32).

Selinexor (KPT-330) is an orally bioavailable, first-in-class selective inhibitor that binds to Cys528 of XPO1, thereby blocking XPO1-dependent nuclear export and promoting the nuclear retention of tumor-suppressor proteins (5, 25, 33–36). It is an FDA approved drug for multiple myeloma (33), and has also shown antitumor activity across diverse cancer cell-line and animal models (5, 7, 25, 29, 37). Currently, it is being evaluated as monotherapy and in combination regimens in clinical trials across a broad range of hematological malignancies and solid tumors (5, 7, 25, 29, 37). Although XPO1 dysregulation and the therapeutic activity of selinexor have been extensively investigated in several hematological malignancies and solid tumors, their relevance in TGCTs remains largely unknown. Here, we profiled XPO1 expression and protein abundance/level in normal testicular tissue, TGCT tissues and TGCT cell lines, and investigated the effects of selinexor in TGCT cell lines. We assessed cell viability, cell-cycle distribution and apoptosis, and evaluated protein levels of p53 and p21 together with their subcellular localization following Selinexor treatment. Our findings indicate that inhibition of XPO1 might be a potential therapeutic strategy in TGCTs.

## 2. MATERIALS AND METHODS

### 2.1. Immunohistochemistry on tissue microarrays

Immunohistochemical staining for XPO1 was performed on tissue microarray (TMA) sections using the BenchMark Ultra automated staining platform (Roche Diagnostics, Germany) as previously described (38). XPO1 antibody (1:50; Santa Cruz Biotechnology, #sc-74454) was used; all other staining reagents were obtained from Medac (Wedel, Germany). Sections were counterstained with Mayer’s hematoxylin (Merck, Darmstadt, Germany). Normal testicular tissue samples (n = 9) were collected, and a TMA was constructed from surgical specimens obtained from a clinically annotated cohort of 148 patients with TGCTs treated with curative or palliative intent at University Hospital Bonn. All diagnoses were confirmed at the Institute of Pathology, University Hospital Bonn. The study was approved by the Institutional Review Board of the University of Bonn (approval no. 187/16). XPO1 immunoreactivity was assessed by semi-quantitative scoring in TMA and normal testicular tissue.

### 2.2. Illumina HT-12 v4 expression microarray analysis of TGCT cell lines and Affymetrix expression microarray analysis of TGCT tissues

Publicly available Illumina HumanHT-12 v4 expression microarray data (GEO accession no. GSE71239) (39) were analyzed to assess XPO1 expression in TGCT cell lines: the seminoma cell line TCam2 (n = 5); the embryonal carcinoma cell lines 2102EP (n = 5) and NCCIT (n=4); and the choriocarcinoma cell line JAR (n = 2) as well as in control fibroblasts MPAF (n=4).

The Affymetrix gene-expression microarray dataset generated by Eckert et al. (40) was used for a meta-analysis to assess XPO1 expression in normal testis (n = 4), GCNIS (n = 3), seminoma (n = 4), embryonal carcinoma (n = 3), teratoma (n = 3), and mixed non-seminoma samples (n = 4).

### 2.3. Cell lines and culture conditions

Human TGCT cell lines representing seminoma (TCam2), embryonal carcinoma (NT2/D1, 2102EP and NCCIT) and choriocarcinoma (JAR and JEG3) were used in this study (38, 39, 41, 42). Human adult fibroblasts cell line MPAF was included as a control (38, 39, 41, 42). Cell line sources, culture-medium formulations and supplements have been described previously (43). All cell lines were maintained at 37 °C in a humidified atmosphere containing 7.5% CO₂. Cell lines were routinely tested for mycoplasma contamination using the TransDetect® PCR Mycoplasma Detection Kit (TransGene Biotech, #FM311-01) and confirmed to be mycoplasma-free before experimental use.

### 2.4. Cell viability assay

Cells were seeded in 96-well plates at 3×10^3^ cells per well and treated with selinexor (MedChemExpress; # HY-17536) for 24, 48 or 72 h. For the 72-h treatment condition, the medium was replenished with fresh selinexor-containing medium after 48 h. Cell viability was determined using the XTT assay as described previously (38, 39, 41, 42). IC_50_ values were determined by fitting concentration-response curves using a four-parameter variable-slope nonlinear regression model (“inhibitor versus response”) in GraphPad Prism.

### 2.5. Cell cycle analysis

Cell-cycle profiles were determined by propidium iodide (PI; BioLegend, #421301) staining as described previously (43). Cells were seeded at 1.5×10^5^ cells per well in six-well plates and treated with selinexor (100 or 200 nM) for 24 h. Cells were harvested, fixed in 70% ethanol, and stained with PI/RNase A, and analyzed using a BD FACSCanto II flow cytometer at the Flow Cytometry Core Facility, University Hospital Bonn and data were analyzed in FlowJo v10.10.

### 2.6. Apoptosis assay

Apoptosis was assessed using the FITC Annexin V Apoptosis Detection Kit with 7-AAD (BioLegend, #640922) as previously described (43). Cells were seeded at 1.5×10^5^ cells per well in six-well plates and treated with selinexor (100 or 200 nM) for 24 or 48 h. Following treatment, cells were washed twice with cold cell staining buffer (BioLegend, #420201) and stained with FITC Annexin V and 7-AAD in Annexin V binding buffer for 15 min at room temperature in the dark. Samples were acquired using a BD FACSCanto II flow cytometer at the Flow Cytometry Core Facility, University Hospital Bonn and data were analyzed in FlowJo v10.10.

### 2.7. Protein Extraction, SDS-PAGE and Western Blot

Cells were seeded at a density of 1×10^5^ cells /mL in 10 cm culture dishes and treated with 100 nM selinexor or DMSO for 24 or 48 hours, followed by total protein extraction and Western blot was performed as previously described (38). Briefly, cells were washed with ice-cold PBS and lysed in RIPA buffer on ice for 5 min. Lysates were sonicated, centrifuged at 14,000g for 15 min at 4°C, and supernatants were collected for protein quantification by BCA assay. Equal amounts of protein (25 µg) were denatured, separated by SDS-PAGE and transferred to 0.45-µm PVDF membranes by wet transfer at 100 V for 1 h at 4 °C. Membranes were blocked for 1 h at room temperature with gentle agitation using the appropriate blocking solution, which was also used to dilute the primary antibodies. Membranes were then incubated with primary antibodies against XPO1 (1:250 in 3% BSA; Santa Cruz Biotechnology, # sc-74454), p53 (1:1000 in 5% BSA; Cell Signaling Technology, # 9282), p21 (1:1000 in 5% BSA; Cell Signaling Technology, # 2947]), and α-tubulin (1:1000 in 5% BSA; Abcam, #ab7291) overnight at 4°C. Subsequently, membranes were incubated with (HRP-conjugated secondary antibodies: anti-mouse IgG (1:2000 in 5% BSA; Agilent Technologies Dako, #P0260) or anti-rabbit IgG (1:2000 in 5% BSA; Agilent Technologies Dako, # P0448) for 1 h at room temperature. Protein signals were detected using chemiluminescent substrates (SuperSignal™ West Pico PLUS, Thermo Fisher Scientific, USA; or WESTAR NOVA 2.0, Cyanagen, Italy) and acquired using a ChemiDoc MP Imaging System (Bio-Rad Laboratories, USA). Following target-protein detection, membranes were stripped for 20 min in mild stripping buffer (25 mM glycine, 1% SDS, pH 2.0), re-blocked and reprobed for α-tubulin using primary and secondary antibodies at the dilutions indicated above, and subsequently imaged.

### 2.8. Immunofluorescence staining

TGCT cells were seeded on glass coverslips in 24-well plates at a density of 5x 10^4^ cells /ml and treated with 100 nM selinexor or DMSO for 24 h. Fixation, permeabilization, blocking and immunostaining were performed as previously described (38), with minor modifications. Cells were fixed in 4% paraformaldehyde, permeabilized with 0.3% Triton X-100, and blocked sequentially with normal horse serum for 20 min and followed by in 5% BSA for approximately 30 min at room temperature. Primary antibodies were diluted in 1% BSA, applied at 100 µl per well and incubated overnight at 4 °C with gentle shaking. The following primary antibodies were used: p53 (1:250; Cell Signaling Technology, # 9282), p21 (1:100; Cell Signaling Technology, # 2947) and α-tubulin (1:1000; Abcam, # ab7291). After washing, coverslips were incubated for 1 h at room temperature fluorophore-conjugated secondary antibodies-the VectaFluor Duet Double Labelling Kit containing DyLight 594 anti-rabbit IgG and DyLight 488 anti-mouse IgG (Biozol, #DK-8828). One drop of the respective secondary-antibody reagent was applied per coverslip. Nuclei were counterstained with Hoechst 33258 (bisbenzimide pentahydrate; Thermo Fisher Scientific) diluted 1:50 in PBS for 20 min at room temperature. Coverslips were mounted using Fluoroshield mounting medium (Sigma-Aldrich, #F6182). Images were acquired using a VisiScope CSU-W1 spinning-disk confocal microscope (Visitron Systems, Germany) at the Core Facility Microscopy, University of Bonn, using VisiView software. Image processing and channel merging were performed in Fiji (ImageJ). Nuclear regions of interest were segmented based on Hoechst nuclear staining following Gaussian blurring and thresholding. Mean fluorescence intensity within nuclear region of interest was quantified in the red channel. For p53 and p21 quantification, 144 - 371 cells were analyzed per condition.

### 2.9. Figure preparation and statistical analysis

Graphs were generated using GraphPad Prism v.10.6.1 and assembled in Inkscape v.1.4.3. Data presented as mean ± SD from independent biological replicates. Statistical comparisons between two groups were performed using unpaired two-tailed Student’s t-tests. Significance levels are indicated as follows: ns, not significant; *P < 0.05; **P < 0.001; ***P < 0.0001; ****P < 0.00001.

## 3. RESULTS

### 3.1. XPO1 is expressed in normal testicular tissue as well as in TGCT tissues and cell lines

As XPO1 is expressed in both normal and malignant cells across multiple organs, we first examined its expression across cell types in the normal human testis using the Human Testis Atlas (44). XPO1 was highly expressed in early germ cells, including spermatogonial stem cells, differentiating spermatogonia, and early and late primary spermatocytes (Supplementary figure 1). Besides, XPO1 expression was also detected in somatic cell populations, including Leydig cells, Sertoli cells, myoid cells and endothelial cells, as well as in macrophages (Supplementary figure 1), suggesting that XPO1 may contribute to both germ cell biology and the maintenance of testicular somatic and immune-cell functions.

Next, a tissue microarray comprising normal spermatogenesis (n = 9), GCNIS (n = 51), seminoma (n = 70) and embryonal carcinoma (n = 27), was analyzed (Figure 1B). Semiquantitative XPO1 immunohistochemical scoring (0 = negative, 1 = weak, 2 = moderate, and 3 = high) was performed to assess XPO1 protein levels and distribution across these tissue types (Figure 1A). High XPO1 staining intensity was observed in all normal testicular tissues and in 98% of GCNIS samples (Figure 1B). In contrast, seminoma and embryonal carcinoma samples showed heterogeneous XPO1 staining intensity, ranging from negative to high (Figure 1B), suggesting increased variability in XPO1 expression following tumor invasion.

**Figure 1:**
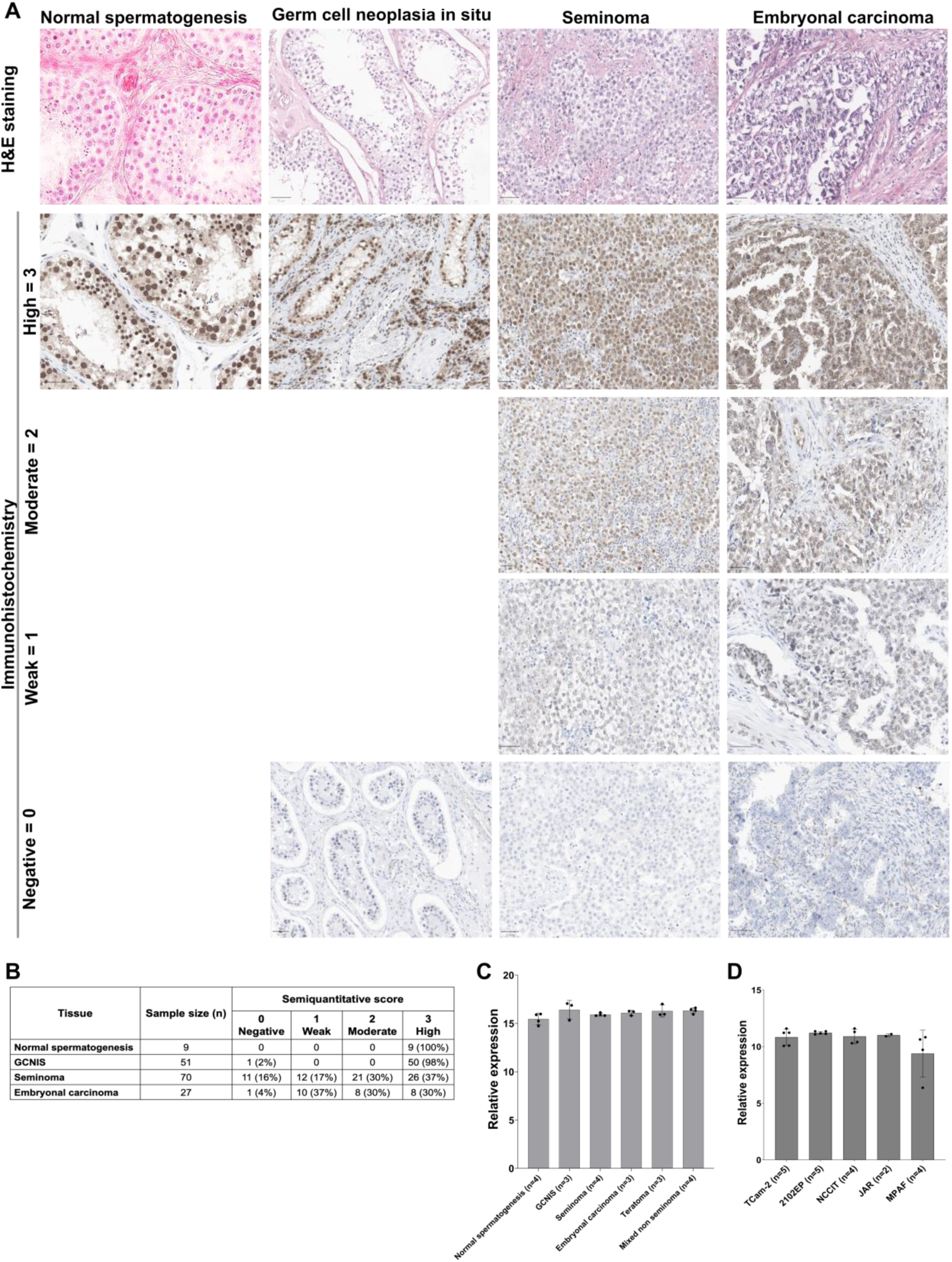
XPO1 expression across normal testis, TGCT tissues and cell lines. **(A)** Representative immunohistochemical staining of XPO1 in normal testis, germ cell neoplasia in situ (GCNIS), seminoma, and embryonal carcinoma from a tissue microarray (TMA), along with corresponding semiquantitative scoring of staining intensity. Scale bars, 50 µm. **(B)** Semi-quantitative assessment of XPO1 immunostaining intensity in normal testicular tissue with intact spermatogenesis (n = 9) and in tissue microarray samples representing GCNIS (n = 51), seminoma (n = 70), and embryonal carcinoma (n = 27). **(C)** Meta-analysis of Affymetrix microarray datasets showing XPO1 expression across normal testis (n = 4), GCNIS (n = 3), seminoma (n = 4), embryonal carcinoma (n = 3), teratoma (n = 3), and mixed non-seminoma samples (n = 4). **(D)** Meta-analysis of Illumina microarray datasets depicting XPO1 expression in TGCT cell lines-TCam2 (n = 5), 2102EP (n = 5), NCCIT (n = 5) and JAR (n = 2) and MPAF fibroblasts (n = 4).

Meta-analysis of Affymetrix microarray datasets comprising normal spermatogenesis, GCNIS, seminoma, embryonal carcinoma, teratoma, and mixed non-seminoma revealed broadly comparable XPO1 transcript levels across the analyzed samples, with relatively lower expression in normal spermatogenic tissue (Figure 1C). Furthermore, meta-analysis of Illumina microarray datasets comprising the TGCT cell lines TCam2, 2102EP, NCCIT and JAR, together with MPAF control fibroblasts, showed that XPO1 expression was broadly comparable across the TGCT cell lines, with relatively higher expression in 2102EP cells (Figure 1D). By contrast, XPO1 expression was lower in MPAF than in all TGCT cell lines, suggesting an association between XPO1 and TGCTs.

Together, these analyses revealed heterogeneous XPO1 expression at the transcript and protein levels across normal testicular tissue, TGCT specimens and TGCT cell lines. As XPO1 contributes to normal cellular homeostasis and is frequently dysregulated in cancer (5–8, 22, 25, 45), we next investigated the effects XPO1 inhibition with selinexor on cell viability, cell cycle distribution and apoptosis in TGCT cell lines.

### 3.2. Targeting XPO1 with selinexor reduces viability of TGCT cell lines

To investigate the effects of selinexor on the viability of TGCT cell lines and control fibroblasts, XTT assays were performed following treatment with increasing selinexor concentrations (50, 100, 200, and 500 nM) for 24, 48, and 72 h. Reduced cell viability was already observed after 24 h of selinexor treatment which was further decreased after 48 h and 72 h in TGCT cells (Figure 2A-F). A dose- and time-dependent reduction of cell viability was observed in TGCT cells (Figure 2A-F; Supplementary figure 2A-F), however, the cell viability of NT2/D1 treated with lowest concentration was slightly higher at 72 h than at 48 h (Figure 2C). Based on IC_50_ values, selinexor potency was categorized as very high (<100 nM), high (>100-200 nM), moderate (>200-500 nM), or low (>500 nM) (Table 1). Following 48 h of selinexor treatment, NT2/D1, 2102EP and JAR cells showed very high sensitivity (IC_50_ <100 nM) (Figure 2B, C, E; Supplementary figure 2B, C, E; Table 1), whereas JEG3 cells were highly sensitive (IC_50_, 108.3 nM) (Figure 2F; Supplementary figure 2F; Table 1) and NCCIT and TCam2 cells displayed moderate sensitivity (IC_50_, 200-500 nM) (Figure 2A, D; Supplementary figure 2A, D; Table 1). Prolonging treatment to 72 h increased sensitivity across all TGCT cell lines: JAR cells were the most sensitive (IC_50_, 4.132 nM), JEG3 and 2102EP cells showed very high sensitivity (IC50, (<100 nM) and NCCIT, NT2/D1 and TCam2 cells were highly sensitive (IC_50_, >100-200 nM)) (Figure 2 A-F; Supplementary figure 2 A-F; Table 1). In contrast, control MPAF showed minimal response even at the highest selinexor concentration with the highest IC₅₀ values at all the time points among all tested cell lines (Figure 2G; Table 1; Supplementary figure 2G). The preferential sensitivity of TGCT cells to selinexor relative to control fibroblast supports a potential therapeutic window for XPO1 inhibition in vitro.

**Figure 2:**
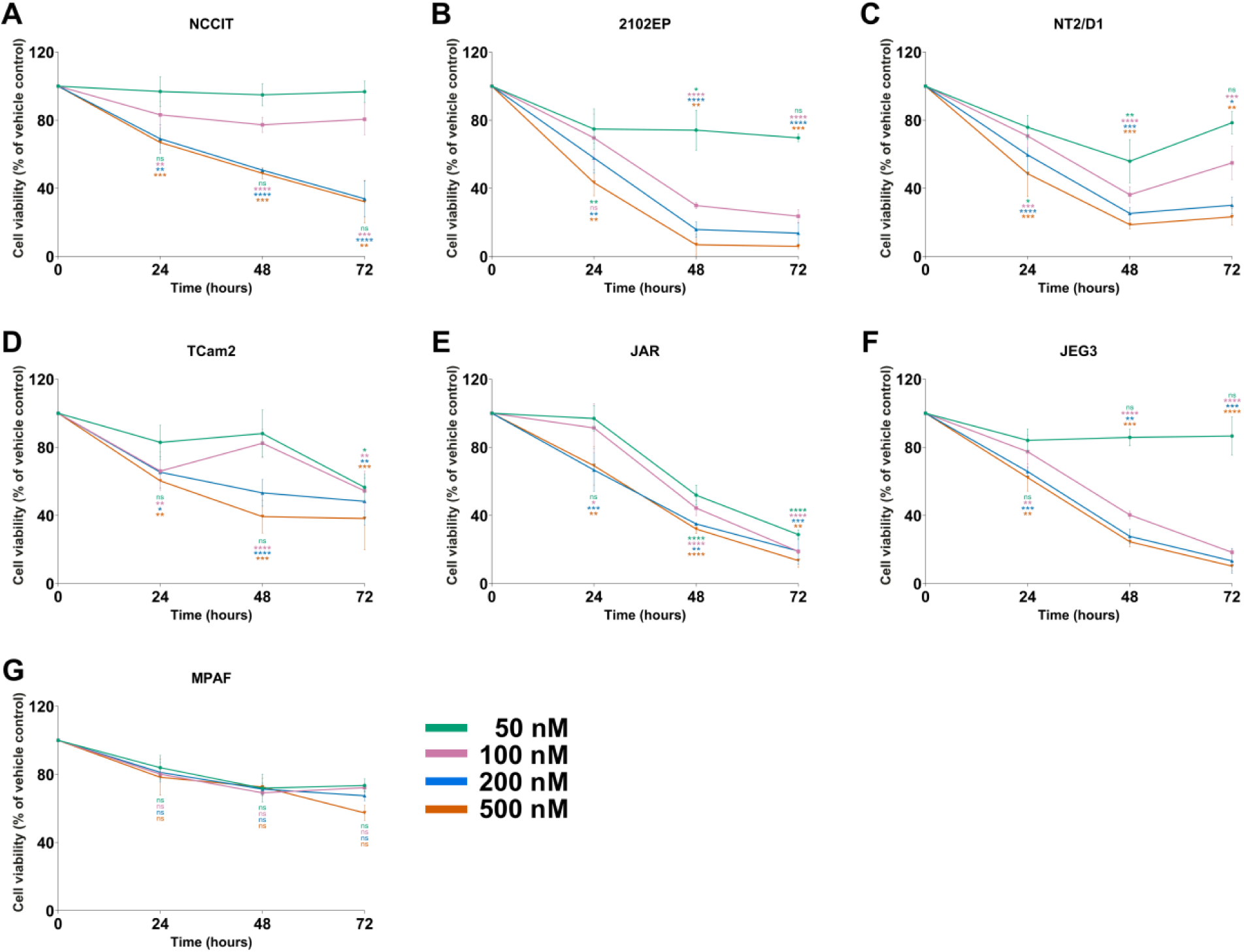
Selinexor reduces TGCT cell viability. XTT viability assays were performed in TGCT cell lines (**A-F**; n = 6) and MPAF fibroblasts (**G**; n = 3) after treatment with selinexor (50, 100, 200, or 500 nM) for 24, 48, or 72 h. DMSO vehicle controls were included for each condition and set to 100% viability; data were normalized to the corresponding vehicle controls and are presented as mean ± SD. Statistical significance was assessed using a two tailed Student’s t test. Asterisks indicate statistical (*P < 0.05, **P < 0.01, ***P < 0.001, ****P < 0.0001), and non-significant differences are denoted as ns (P > 0.05). Asterisk color indicates the selinexor concentration.

**Table 1:** IC50 values (nM) for TGCT and control cell lines following Selinexor treatment, determined using the XTT viability assay. Results are presented for 24, 48, and 72 h post-treatment. Color coding denotes potency: red (0-100 nM, very high potency), blue (>100-200 nM, high potency), Yellow (>200-500 nM, moderate potency), and Teal (>500 nM, low potency).

| Entity | Cell line | IC50 (nM) |  |  |
| --- | --- | --- | --- | --- |
|  |  | 24 h | 48 h | 72 h |
| Embryonal carcinoma | NT2/D1 | 444.8 | 58.66 | 123.9 |
|  | 2102EP | 331.7 | 75.55 | 67.83 |
|  | NCCIT | >500 | 352.9 | 198.2 |
| Seminoma | TCam2 | >500 | 296.4 | 138.0 |
| Choriocarcinoma | JAR | >500 | 54.96 | 4.132 |
|  | JEG3 | >500 | 108.3 | 73.81 |
| Fibroblast | MPAF | >500 | >500 | >500 |

### 3.3. Selinexor induces cell cycle arrest and apoptosis in TGCT cells

To determine whether selinexor-induced reductions in cell viability were associated with altered cell-cycle progression, TGCT cell lines and control fibroblasts were treated with 100 or 200 nM selinexor for 24 h and subsequently analyzed by flow cytometry. TGCT cell lines exhibited heterogeneous, dose-dependent cell-cycle responses to selinexor, with accumulation in either the G1 or G2/M phase (Figure 3A-F). Selinexor induced a dose-dependent increase in the NCCIT, 2102EP and TCam2 cells in the G1 phase (Figure 3A, B, D), whereas NT2/D1, JAR and JEG3 cells accumulated in the G2/M phase (Figure 3C, E, F). In MPAF, selinexor elicited only a modest increase in the G1 phase under the same conditions (Figure 3G), markedly less pronounced than the cell-cycle changes observed in TGCT cell lines.

**Figure 3:**
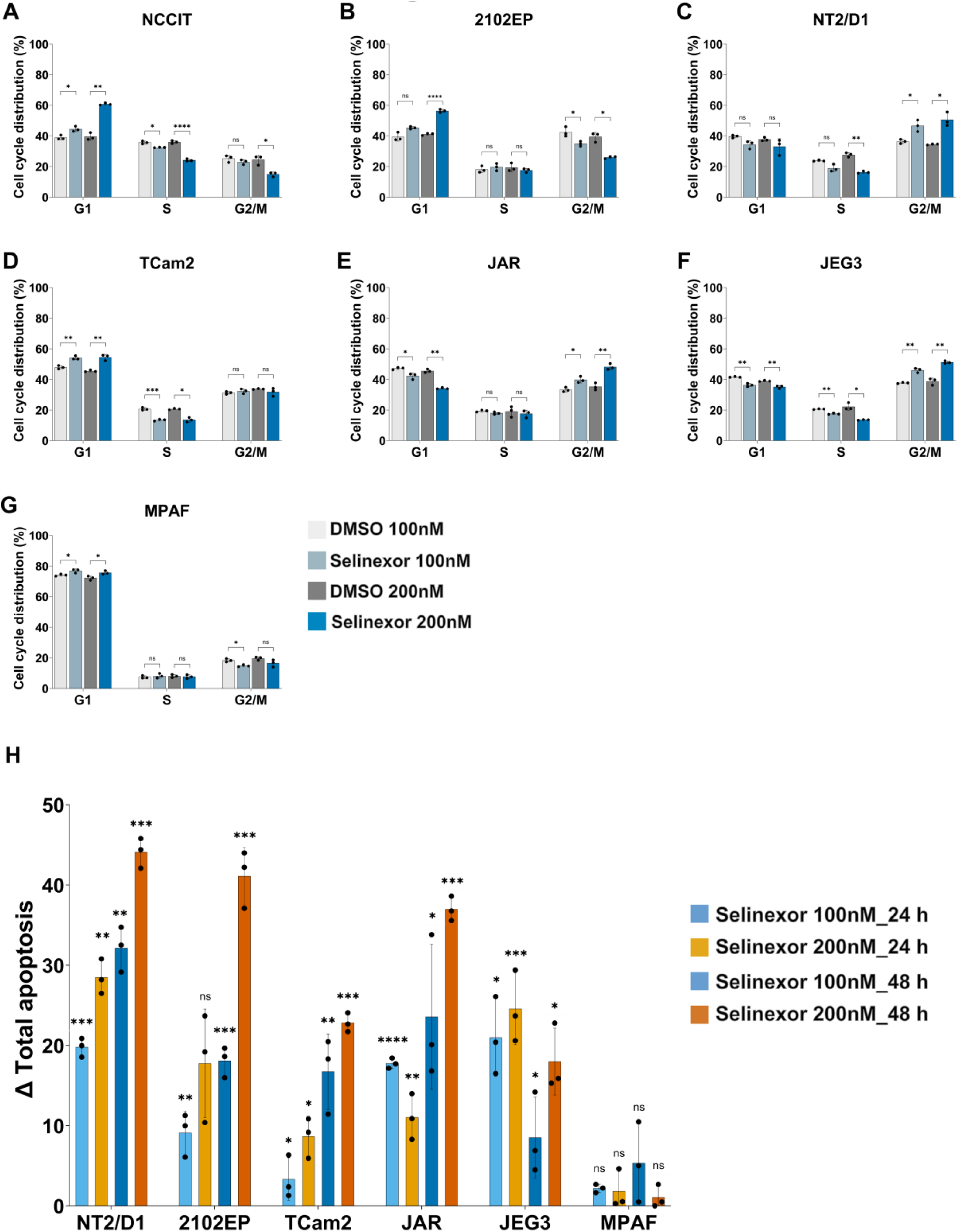
Selinexor induces cell-cycle arrest and apoptosis in TGCT cell lines. Cell-cycle distribution of TGCT cell lines (**A-F**) and control fibroblast MPAF (**G**) were treated with Selinexor (n = 3, data are shown as mean ± SD). TCam2, 2102EP, NCCIT, NT2/D1, JAR and JEG3 cells, and MPAF fibroblasts, were treated with selinexor (100 or 200 nM) or vehicle control (DMSO) for 24 h. Cells were stained with propidium iodide and analysed by flow cytometry. Selinexor increased the G1-phase fraction in NCCIT, 2102EP and TCam2 cells, whereas NT2/D1, JAR and JEG3 cells showed accumulation in G2/M phase. MPAF fibroblasts showed a modest increase in the G2/M fraction. **H.** Quantification of selinexor-induced apoptosis. TGCT cell lines and MPAF fibroblasts were treated with selinexor (100 or 200 nM) or DMSO for 24 or 48 h and analyzed by Annexin V-FITC and 7-AAD flow cytometry (n = 3, data are shown as mean ± SD). Δ total apoptosis denotes the difference in total apoptotic cells between selinexor-treated and DMSO-treated samples and is expressed in percentage points. Selinexor increased apoptosis in TGCT cell lines in a dose- and time-dependent manner, whereas only minimum apoptosis was observed in MPAF fibroblasts. Statistical analysis was performed using a two-tailed Student’s t-test relative to DMSO (*P < 0.05, **P < 0.01, ***P < 0.001, ****P < 0.0001; ns, not significant (P > 0.05).

Given the observed selinexor-induced reduced cell viability and alterations in cell-cycle distribution, we next assessed the apoptotic response to selinexor in TGCT cells. TGCT cell lines and control MPAF were treated with 100 or 200 nM selinexor for 24 or 48 h, and apoptosis was quantified by flow cytometry following FITC-Annexin V/7-AAD staining. Selinexor induced dose- and time-dependent apoptosis across all TGCT cell lines, as determined by an increased proportion of total apoptotic cells relative to DMSO-treated vehicle controls (Figure 3H). The apoptotic response was most pronounced in NT2/D1 and 2102EP cells, followed by JAR and JEG3 cells, whereas TCam2 cells showed a more modest, yet significant, increase in apoptosis (Figure 3H). By contrast, selinexor elicited only minimal apoptosis in MPAF under the same conditions (Figure 3H). Together, these data show that selinexor selectively alters cell-cycle distribution and induces apoptosis in TGCT cells.

### 3.4. Selinexor modulates p53 and p21 levels and localization in TGCT cells

To investigate mechanisms underlying the selinexor response, we examined p53 and its downstream effector p21, key regulators of cell-cycle progression and apoptosis that are subject to XPO1-mediated nuclear export. TGCT cells were treated with 100 nM selinexor for 24 or 48 h, and XPO1, p53 and p21 protein levels were assessed by western blotting using α-tubulin as a loading control (Figure 4A-E). Uncropped western blot images for Figure 4A-E are provided in Supplementary Figure 3A-C. Selinexor differentially affected XPO1 protein levels across TGCT cell lines (Figure 4A-E). XPO1 levels were reduced in the choriocarcinoma cell lines JAR and JEG3 (Figure 4C-D) but remained largely unchanged in the other TGCT cell lines (Figure 4A, B, E), demonstrating a cell-line-specific effect of selinexor on total XPO1 abundance. Despite the heterogeneous effects of selinexor on total XPO1 protein levels, selinexor caused increased p53 and p21 protein levels in all TGCT cell lines (Figure A-E). These increases were evident at 24 h and persisted at 48 h, indicating a sustained p53 and p21 response to selinexor across the TGCT cell lines.

**Figure 4:**
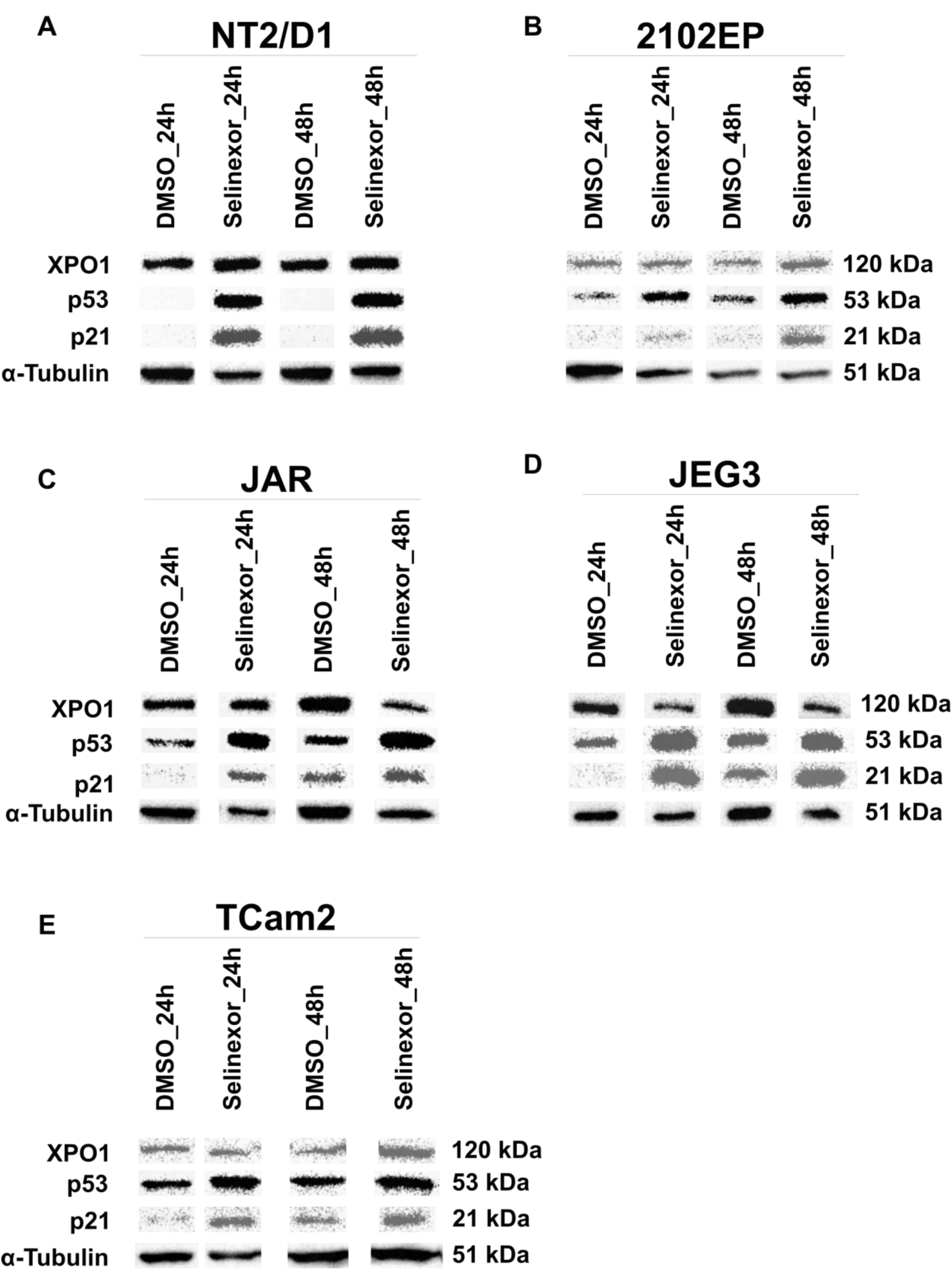
Selinexor modulates XPO1, p53 and p21 protein abundance in TGCT cell lines. Immunoblot analysis of XPO1, p53 and p21 in NT2/D1 (**A**), 2102EP (**B**), JAR (**C**), JEG3 (**D**) and TCam2 (**E**) cells following treatment with 100 nM selinexor for 24 or 48 h. XPO1 protein abundance was reduced or remained unchanged following selinexor treatment, whereas p53 and p21 protein abundance increased across the indicated TGCT cell lines.

Building on the sustained increase in total p53 and p21 protein levels, we next assessed their subcellular distribution in NT2/D1, TCam2 and JAR following treatment with 100 nM selinexor for 24 h. Immunofluorescence analysis revealed an apparent increase in nuclear p53 and p21 staining in NT2/D1, TCam2 and JAR relative to DMSO-treated vehicle controls (Figure 5A-C, Supplementary figure 4A-C), indicating that selinexor alters the abundance and subcellular distribution of these proteins in TGCT cells.

**Figure 5:**
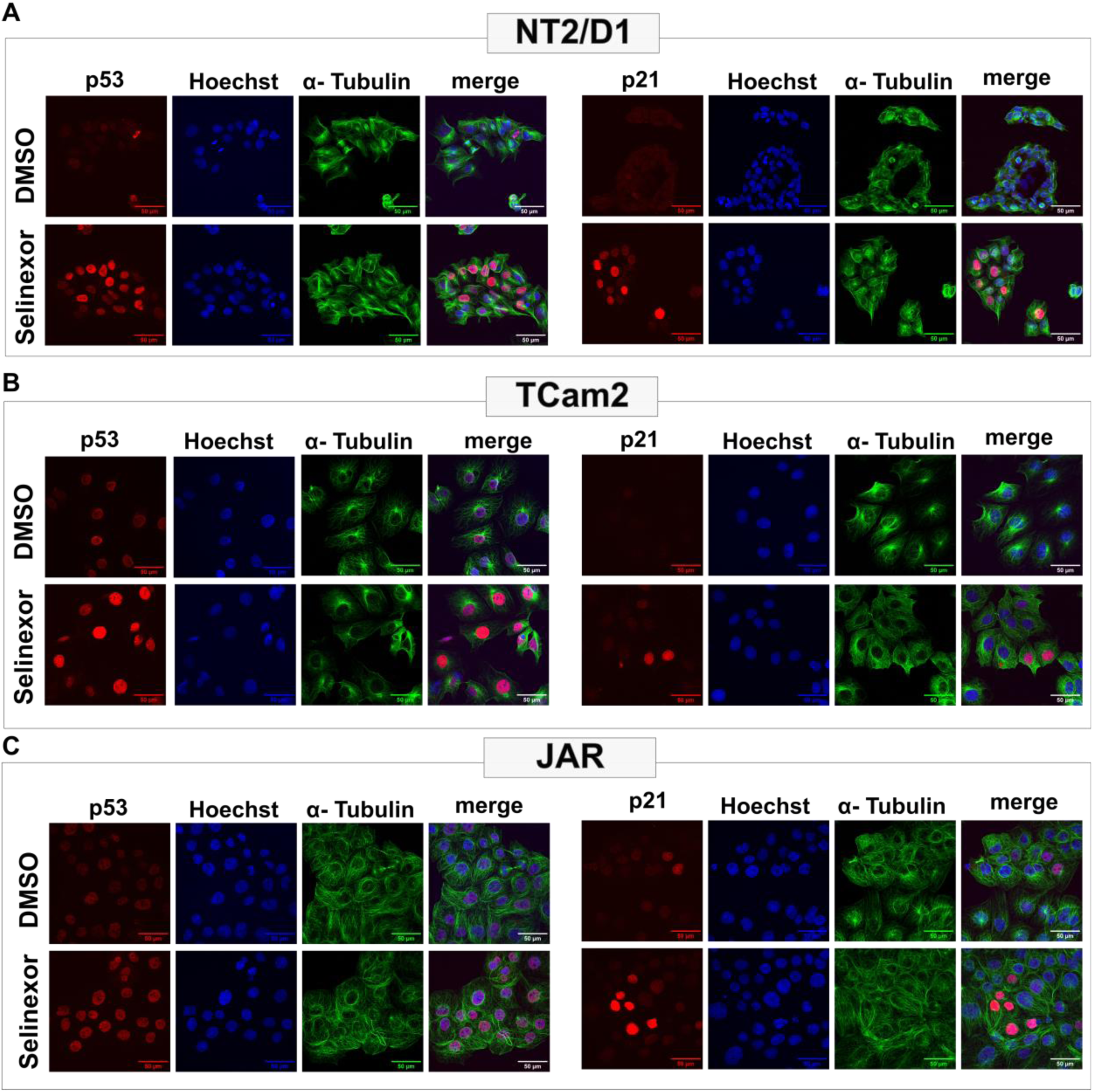
Selinexor promotes nuclear accumulation of p53 and p21 in TGCT cell lines. Immunofluorescence analysis of p53 and p21 localization in NT2/D1 (**A**), TCam2 (**B**) and JAR (**C**) cells following treatment with 100 nM selinexor for 24 h. Selinexor increased nuclear p53 and p21 immunoreactivity in all the cell lines. p53 or p21, red; α-tubulin, green; nuclei, Hoechst (blue). Scale bar, 50 μm.

## 4. DISCUSSION

Our findings demonstrated XPO1 as a potential therapeutic target in TGCTs. Targeting XPO1 with selinexor reduced TGCT cell viability, induced cell-cycle accumulation in the G1 or G2/M phases, and increased apoptosis. By contrast, selinexor exerted limited effects in control fibroblasts under the conditions tested, suggesting differential sensitivity between malignant and non-malignant cells. Furthermore, selinexor increased total p53 and p21 protein levels and their nuclear accumulation in TGCT cell lines. As p53 and p21 are central regulators of cell-cycle arrest and apoptosis, their nuclear retention may contribute to the antitumor effects of XPO1 inhibition in TGCT cells.

Immunohistochemical analysis revealed more intense XPO1 expression in normal testicular tissue than in GCNIS, seminoma and embryonal carcinoma, indicating that XPO1 is not selectively elevated in TGCTs. Consistent with its broad expression in early germ-cell and somatic-cell populations in the adult human testis transcriptional cell atlas, XPO1 may have functions in normal testicular biology. Although the specific role of XPO1 in male germ cells remains unknown, studies in porcine oocytes suggest that XPO1-mediated nuclear export contributes to germinal-vesicle maintenance and meiotic resumption during oocyte maturation, supporting a potential role for XPO1 in germ-cell and reproductive biology (46). Noteworthy, in murine spermatogonia and spermatocyte cell lines, selinexor at relatively high concentrations (IC_50_, 6.85-16.73 µM) reduced cell viability and proliferation, induced apoptosis, impaired migration, and increased DNA damage and cellular stress (47).

Despite heterogeneous XPO1 expression across TGCT specimens, we investigated the antitumor activity of selinexor in TGCT cell lines, as the consequences of XPO1 activity may be context dependent (5–7, 22, 25). While XPO1-mediated nuclear export supports cellular homeostasis in normal cells, its aberrant activity in cancer can promote the cytoplasmic mis-localization and functional inactivation of tumor suppressors and cell-cycle regulators (5–7, 22, 25, 45).

Following 72 h of exposure, selinexor showed potent activity across TGCT cell lines, with IC_50_ values ranging from 4.1 to 198 nM. This range is comparable to that reported in selinexor-sensitive cell lines of triple-negative breast cancer (11 - 550 nM) (48), T-cell lymphoblastic lymphoma (18.70 - 46.87 nM) (49), mantle-cell lymphoma (44.06 - 239.2 nM) (14), multiple myeloma (100 - 350 nM) (50), sarcoma (50 - 2500 nM in TP53-wild-type, 250 - 500 nM in TP53-mutant and 100 - 1500 nM in TP53-null cell lines) (51), glioblastoma (85 - 550 nM) (52), bladder cancer (210-440 nM) (53), and non-small-cell lung cancer (25-995 nM) (54), supporting further preclinical evaluation of XPO1 inhibition in TGCT. By contrast, the control fibroblasts exhibited a substantially higher IC_50_ (1432 nM) at 72h, indicating preferential in vitro sensitivity of TGCT cells to selinexor.

Selinexor elicited heterogeneous, dose-dependent changes in cell-cycle distribution across TGCT cell lines, with G1 accumulation in NCCIT, 2102EP and TCam2 cells and G2/M accumulation in NT2/D1, JAR and JEG3 cells. These cell-line-specific responses may reflect differences in molecular context and in the relative contribution of XPO1 cargoes that regulate the G1 and G2/M checkpoints in TGCTs. Heterogeneous cell-cycle responses to selinexor have also been reported in hepatocellular carcinoma and thymic epithelial tumors cell lines, with accumulation in both G1 and G2/M phases in one cell line and predominantly G2/M accumulation in another cell line (55, 56). Furthermore, selinexor was also reported to induce cell-line-specific accumulation within a single cell-cycle phase in several cancers. Predominant G1 accumulation was observed in multiple myeloma (57), diffuse large B-cell lymphoma (58), non-small-cell lung cancer (54), triple-negative and other breast cancers (59, 60), gastric cancer (20, 21), cutaneous T-cell lymphoma (61) and gastrointestinal stromal tumors cell lines (62), whereas predominant G2/M accumulation has been reported in thymic epithelial tumors (56) and hepatocellular carcinoma (55) cell lines. Collectively, these observations support a cell-context-dependent effect of selinexor on cell-cycle progression, consistent with the heterogeneous G1 and G2/M responses observed in TGCT cell lines. By contrast, selinexor induced only a modest increase in the G1-phase fraction in MPAF, which was less pronounced than the cell-cycle alterations observed in TGCT cell lines. These findings suggest that, under the conditions tested, TGCT cells are more susceptible than MPAF to selinexor-induced alterations in cell-cycle distribution.

Selinexor induced marked dose- and time-dependent apoptosis across TGCT cell lines, with the strongest effects in embryonal carcinoma and choriocarcinoma cells, consistent with its reported pro-apoptotic activity in mantle cell lymphoma (14, 63), diffuse malignant peritoneal mesothelioma (64), colorectal cancer (24, 65), bladder cancer (66), liposarcoma (67), cutaneous T-cell lymphoma (61), multiple myeloma (68), gastric cancer (20, 21), non-small cell lung cancer (54) and thymic epithelial tumors (56) cell lines. By contrast, MPAF exhibited minimal apoptosis under identical treatment conditions, consistent with a greater apoptotic response to selinexor in TGCT cells in vitro.

Although selinexor consistently reduced cell viability and induced cell-cycle arrest and apoptosis, its effects on total XPO1 protein abundance differed among TGCT cell lines. XPO1 protein levels were reduced in only choriocarcinoma cells but remained largely unchanged in the other TGCT cell lines, indicating that a decrease in total XPO1 protein abundance is not mandatorily required for selinexor-induced reductions in cell viability, cell-cycle arrest and apoptosis. This observation is consistent with the established mechanism of selinexor, which covalently engages Cys528 within the XPO1 cargo-binding pocket to block XPO1-mediated nuclear export of proteins bearing leucine-rich nuclear-export signals (5–7, 25, 29, 35, 45, 64). Thus, in TGCT cells, selinexor activity is consistent with functional XPO1 inhibition rather than depletion of total XPO1 protein, promoting the nuclear retention of tumor-suppressive and growth-regulatory cargoes.

In TGCT cells, selinexor increased total p53 and p21 protein abundance and promoted their nuclear accumulation, consistent with findings reported in gastric cancer (20, 21), cutaneous T-cell lymphoma (61), non-small-cell lung cancer (54), multiple myeloma (36), cholangiocarcinoma (69, 70) and colorectal cancer (24, 71). Given the low (1-5%) frequency of TP53 mutations in TGCTs (72), selinexor-induced nuclear accumulation of p53 may contribute to modulation of the p53-p21 axis and thereby to cell-cycle arrest and apoptosis.

Selinexor demonstrated anti-tumor activity as monotherapy and potential to enhance the efficacy of combination treatments in diverse cancers (5, 6, 8, 10–12, 14, 16, 20, 22, 24, 25, 29–32, 36, 37, 47, 49, 50, 52, 58, 61, 63, 68–71). Furthermore, selinexor may help overcome treatment resistance by enhancing cisplatin sensitivity in germinal-center B-cell-like-DLBCL cells (73), ovarian cancer (74), Small Cell Lung Cancer (75). Although these observations support the potential of selinexor-based combination strategies in refractory malignancies, their relevance to TGCT remains unknown. Given the clinical importance of cisplatin resistance in TGCTs treatment failure, future studies should evaluate selinexor in cisplatin-resistant TGCTs, both as a single agent and in combination with cisplatin. Such studies will be required to determine whether XPO1 targeting enhances platinum-induced cytotoxicity, restores cisplatin sensitivity or exposes alternative therapeutic vulnerabilities in refractory TGCTs. Defining the contribution of additional tumor-suppressive XPO1 cargoes will further clarify the mechanisms underlying selinexor activity. Collectively, these data provide a rationale for continued preclinical evaluation of selinexor, alone and in combination-based strategies for TGCT including cisplatin-resistant disease.

## 5. CONCLUSION

Overall, we demonstrated that the inhibition of XPO1 by selinexor reduced TGCT cell viability and induced cell-cycle arrest and apoptosis, accompanied by nuclear accumulation of p53 and p21. These findings support further preclinical evaluation of selinexor in TGCTs.

## AUTHOR CONTRIBUTIONS

HS and RI designed the study. RI, HS and GK contributed to the conceptual development of the project. RI, AW, LD, AS and VABA performed the meta-analysis of publicly available gene-expression microarray datasets, XTT viability, cell cycle, apoptosis and western blot analyses. IFB performed the TMA analysis. AL assisted with optimization of western blot analysis. AW performed immunofluorescence analyses with substantial support from AK and GEM. RI and HS drafted the manuscript. RI, HS and GK critically revised the manuscript. All authors read and approved the final manuscript.

## ACKNOWLEDGMENTS

We thank Greta Zech and Susanne Steiner for excellent technical assistance. We are grateful to the Flow Cytometry Core Facility of the Institute of Molecular Medicine and Experimental Immunology, University Hospital Bonn for technical support and access to instrumentation. We also acknowledge the Microscopy Core Facility of the Medical Faculty, University of Bonn, for technical support and access to instrumentation funded by the DFG (388171357).

