## Supplemental Information for "XPO1 Inhibition by Selinexor Induces Nuclear p53 and p21 Accumulation, Cell-Cycle Arrest and Apoptosis in Testicular Germ Cell Tumors"

**Supplementary figure 1: XPO1 expression in young adult human testicular tissue.** XPO1 expression data highlighted in red in human testis, extracted from the Human Testis Atlas and adapted from Guo et al. 2018. XPO1 is highly expressed in Spermatogonial stem cells, differentiating spermatogonia, early and late spermatocytes, as well as in macrophages, Sertoli cells, Leydig cells, myoid cells, and endothelial cells.

**Supplementary figure 2: Nonlinear regression analysis of Selinexor sensitivity and IC<sub>50</sub> calculation.** IC<sub>50</sub> values for selinexor at 24, 48, and 72 h were calculated using four-parameter nonlinear regression. The dose-response curves were derived from the XTT cell viability data presented in Figure 2.

**Supplementary figure 3: Raw western blot images after selinexor treatment in TGCT cells.** Uncropped western blot images showing protein levels after 24 h and 48 h of selinexor treatment in TGCT cell lines. (A) NT2/D1 (left) and JAR (right), and (B) 2102EP (left) and TCam2 (right), (C) JEG3.

**Supplementary figure 4: Nuclear p53 and p21 fluorescence intensities following selinexor treatment in TGCT cells.**

Violin plots show the distribution of mean nuclear fluorescence intensities of p53 and p21 in TGCT cells treated with 100 nM selinexor or an equivalent volume of DMSO for 24 h. Centre line indicates the median; dashed lines indicate the quartiles. Statistical significance was determined using Student's t-test (ns: not significant; \*p<0.05; \*\*p<0.01; \*\*\*p<0.001; \*\*\*\*p<0.0001)

### Supplementary figure 1

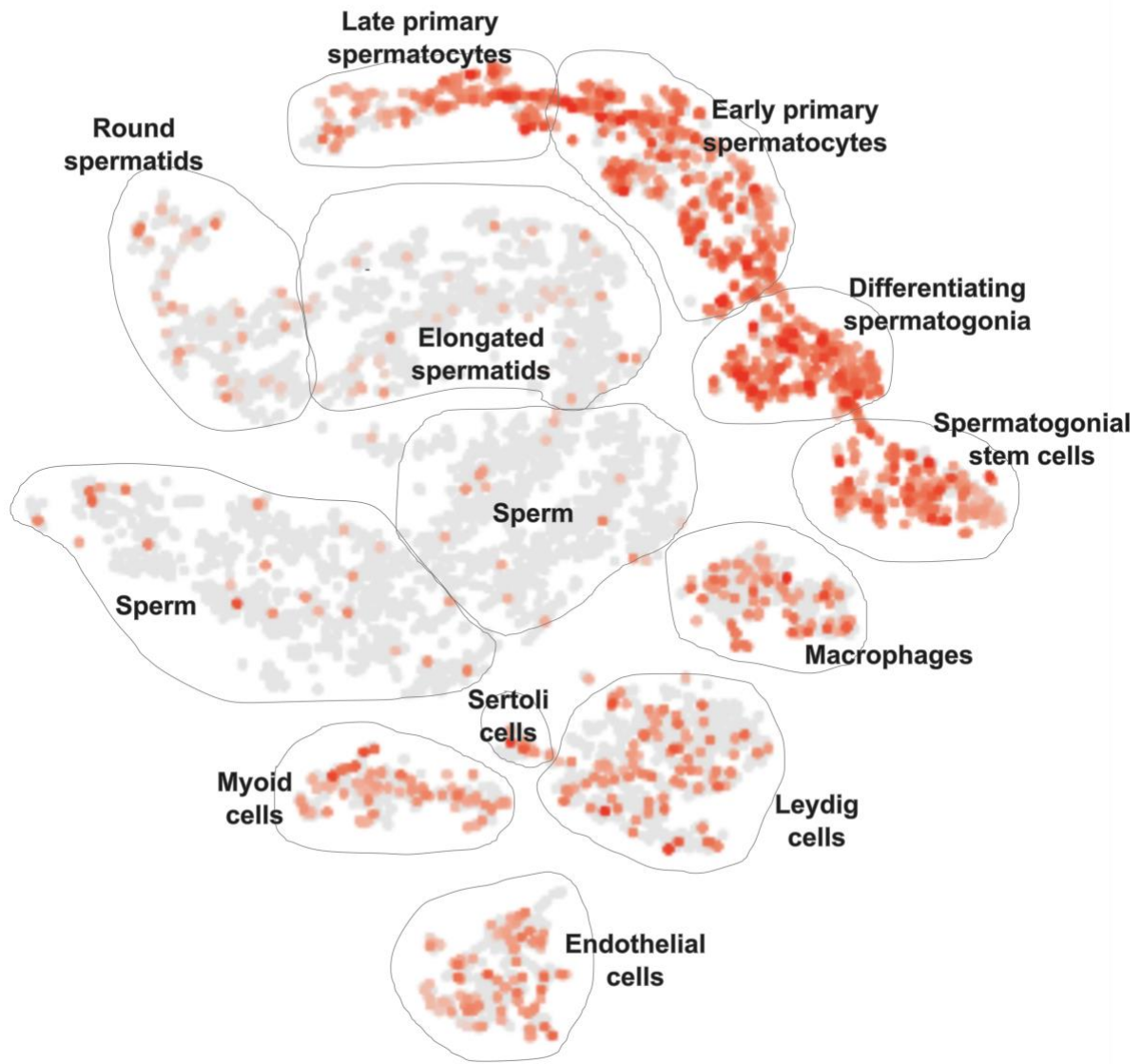

32
33
34
35
36
37
38
39
40

Supplementary figure 2

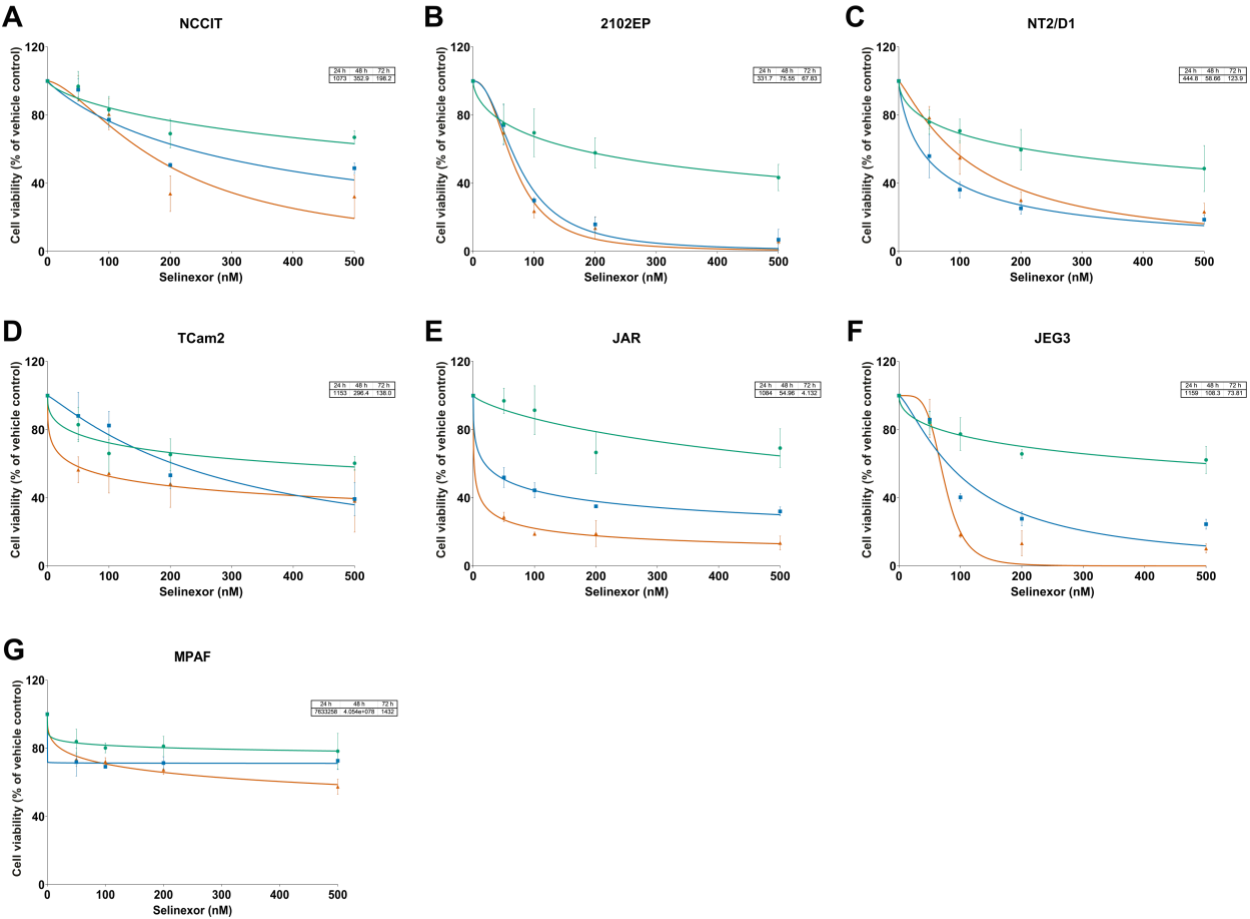

Supplementary figure 3

A

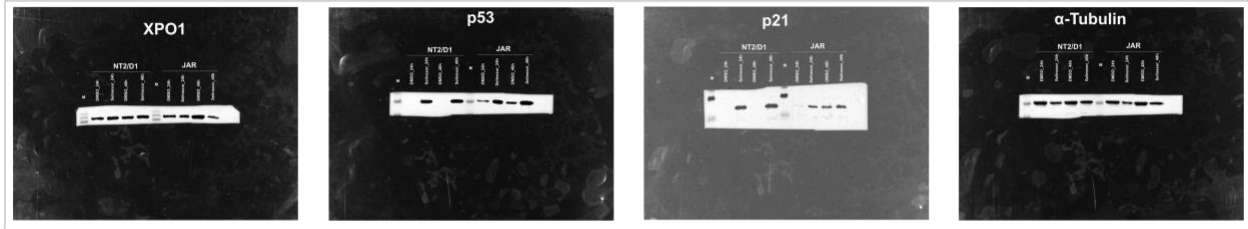

B

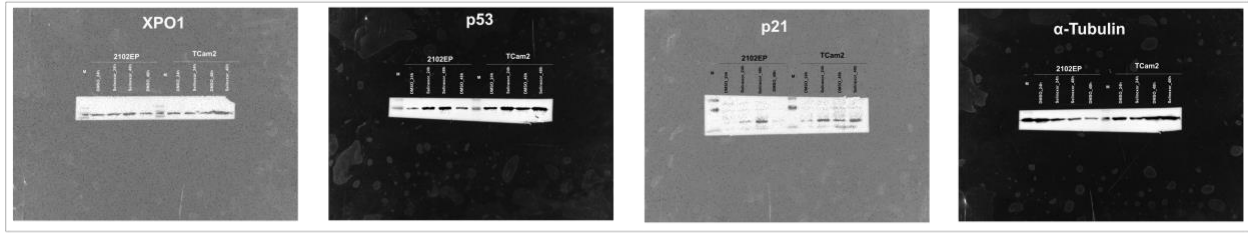

C

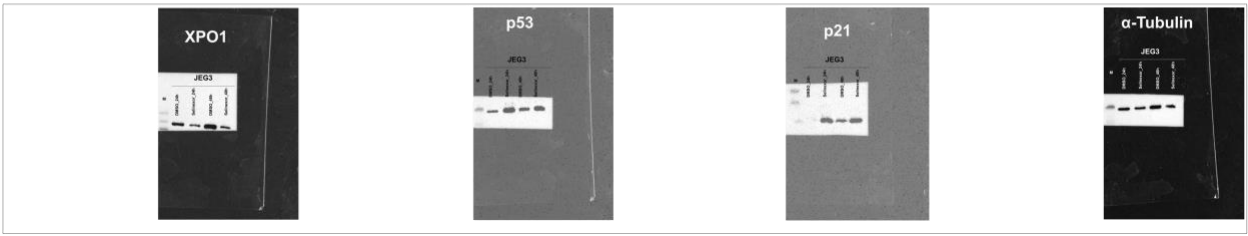

### Supplementary figure 4

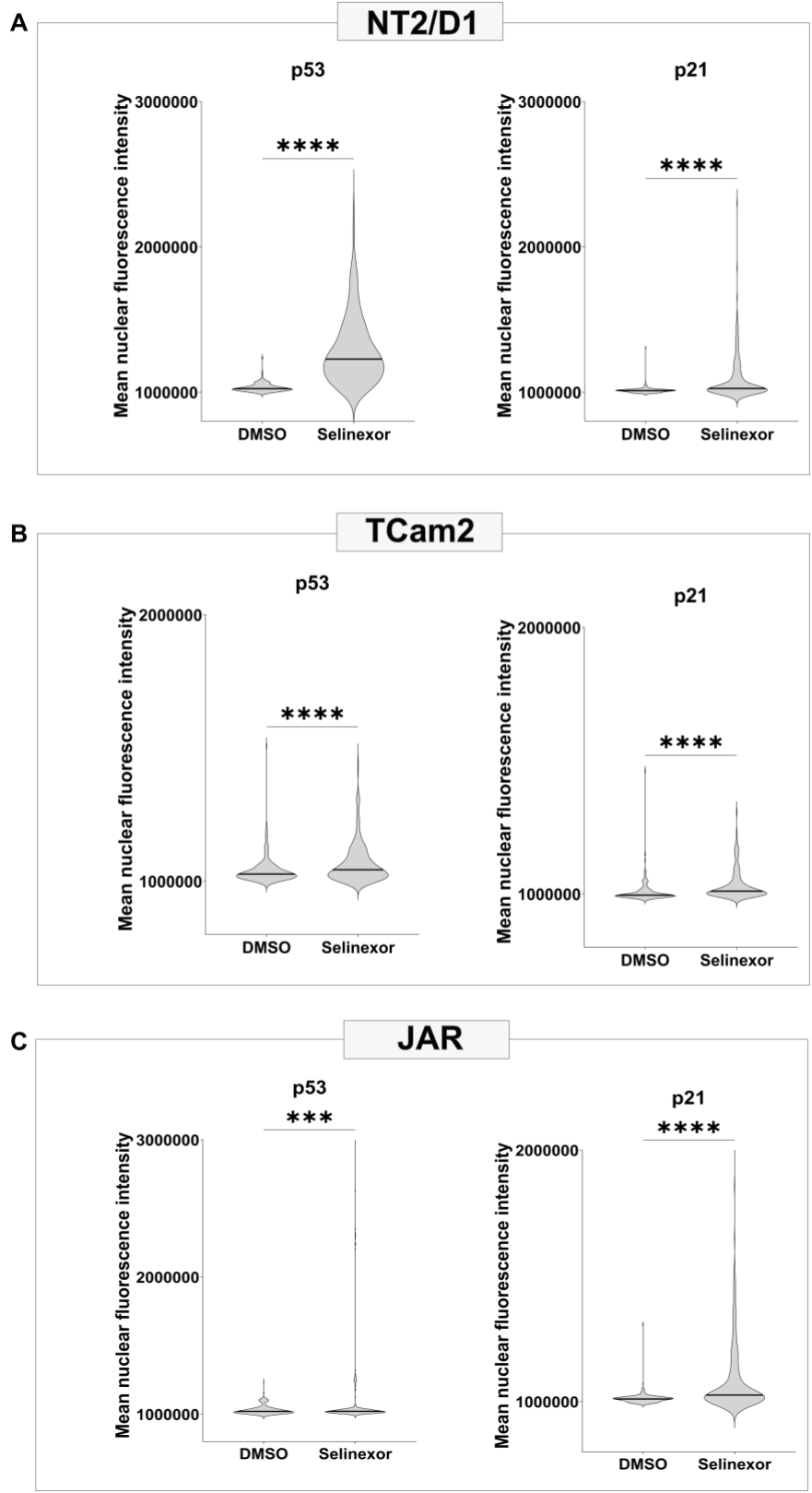
